# CANNABIDIOL MODULATES CLASSICAL AND NON-CLASSICAL HLA EXPRESSION IN HUMAN CHORIOCARCINOMA CELL LINE

**DOI:** 10.64898/2025.11.28.691248

**Authors:** Kevin I. Martínez, María B. Palma, Fernando J. Sepúlveda, Edgardo D. Carosella, Marcela N. García, Fernando L. Riccillo

## Abstract

Cannabidiol (CBD) modulates diverse signaling pathways with potential relevance to tumor immune escape, however its impact on the regulation of classical and non-classical HLA class I molecules remains incompletely understood. Here, we examined the mechanisms by which CBD regulates HLA expression in JEG-3 choriocarcinoma cells, focusing on cannabinoid-related receptors and intracellular Ca²⁺ signaling. CBD increased the expression of classical HLA class I genes—most notably HLA-C—while reducing HLA-G levels, a non-classical HLA class I molecule that acts as a local immunosuppressor. Receptor profiling revealed constitutive expression of CB1 and CB2, whereas GPR55 and PPARγ expression became detectable only after CBD exposure. Functional inhibition assays showed that HLA-G downregulation was selectively attenuated by CB1 blockade, with no meaningful contribution from CB2 or GPR55. In contrast, CBD-induced HLA-C upregulation required GPR55 and CB2 activity, while being unaffected by CB1 inhibition, indicating distinct receptor pathways for classical and non-classical HLA regulation. Calcium chelation using BAPTA further demonstrated that HLA-G modulation is highly sensitive to intracellular Ca²⁺ reduction, whereas classical HLA expression required higher BAPTA concentrations to be affected. Altogether, these findings identify CBD as a dual immunomodulatory agent capable of enhancing tumor immune visibility while limiting immunotolerant HLA-G expression through receptor-specific and Ca²⁺-dependent mechanisms.

## INTRODUCTION

The endocannabinoid system (ECS) plays a fundamental role in regulating multiple physiological processes, including metabolism, cell survival, and immune responses [1–3]. It comprises endogenous ligands such as anandamide (AEA) and 2-arachidonoylglycerol (2-AG), their canonical receptors CB1 and CB2, and associated metabolic enzymes [4–7]. Dysregulation of the ECS has been implicated in various pathological conditions, ranging from neurodegenerative and metabolic disorders to cancer. In addition, phytocannabinoids (the plant-derived cannabinoids from *Cannabis sativa*) exert their effects by mimicking the actions of endogenous cannabinoids, primarily through the CB1 and CB2 receptors. Among them, the two most abundant and therapeutically characterized compounds are Δ9-tetrahydrocannabinol (THC) and cannabidiol (CBD). CBD has very low affinity for CB1/CB2, although it displays pleiotropic activity by engaging alternative targets, including G protein–coupled receptors (GPR55, GPR18), TRP channels (TRPV1, TRPV4), serotonin 5-HT1A, GABA receptors, and PPARγ [8,9].

Cannabinoids have long been used for palliative purposes in oncology, particularly for their analgesic, antiemetic, and orexigenic effects [10–12]. However, increasing evidence indicates that they also exert direct antitumor effects in both *in vitro* and *in vivo* models. THC and CBD have been reported to inhibit proliferation, migration, and angiogenesis, as well as to induce apoptosis in several tumor types, including glioblastoma, breast, prostate, lung, and pancreatic cancers [13–17]. Nevertheless, the underlying mechanisms remain incompletely understood due to their pleiotropic signaling pathways and concentration-dependent effects. Furthermore, CBD offers notable therapeutic advantages. While the clinical use of THC is restricted by its psychotropic activity, CBD has no psychoactive effects and displays a more favorable safety profile, thereby reinforcing its potential as an antitumor and immunomodulatory agent in cancer biology. It should not be overlooked that the combined use of CBD and THC remains a relevant therapeutic possibility, as CBD can modulate THC-mediated signaling through its allosteric effects on CB1 and CB2 [18,19], allowing their contributions to be adjusted and potentially optimized for clinical applications.

Human Leukocyte Antigen (HLA) class I molecules, the human equivalent of MHC-I, are central mediators of immune recognition and play a pivotal role in shaping the balance between immune activation and tolerance. Classical HLA class I molecules (HLA-Ia: HLA-A, -B, -C), broadly expressed on all nucleated cells, are essential for presenting endogenous antigens to cytotoxic T lymphocytes, enabling recognition and elimination of infected or transformed cells. In contrast, non-classical subtypes (HLA-Ib: HLA-E, -F, -G, -H), display a more restricted tissue distribution and function. Among them, the HLA-E and HLA-G, are characterized by tolerogenic functions that are often exploited by viruses and tumors to evade the host immune responses [20]. HLA-G are predominantly expressed at the maternal–fetal interface and contribute to immune tolerance by interacting with inhibitory receptors on NK cells, T and B lymphocytes, and antigen-presenting cells [21, 22]. However, its aberrant expression in tumors has been strongly associated with immune evasion and cancer progression [22–24].

Alterations in HLA expression are increasingly recognized as key determinants of tumor responsiveness to immunotherapies, particularly immune checkpoint inhibitors. Modulation of HLA molecules by tumor cells not only impairs antigen presentation but also promotes a tolerogenic microenvironment, thereby limiting the efficacy of cytotoxic T-cell responses. Recent evidence also suggests that calcium homeostasis may participate in the regulation of HLA expression in tumor cells. In the renal carcinoma cell line named RCC7, stable HLA-G expression was shown to induce an aberrant upregulation of calcium transporters [25]. Moreover, studies in pancreatic ductal adenocarcinoma (PDAC) indicate that dysregulated calcium influx promotes autophagy-mediated lysosomal degradation of MHC class I molecules [26, 27]. Altogether, these observations raise the possibility that calcium-dependent pathways may influence classical and non-classical HLA molecules, although this remains largely unexplored. A better understanding of HLA regulation, including its pharmacological modulation, is therefore essential for developing strategies that enhance antitumor immunity and improve the effectiveness of current immunotherapeutic approaches.

Cannabinoids, through their diverse signaling pathways, have been shown to modulate several of the key molecules involved in tumor immunosurveillance. For instance, there is evidence that THC may impair the efficacy of PD-L1/PD-1 immune checkpoints blockade by acting on CB2 receptors expressed on tumor-specific T cells [28]. In contrast, CBD has been shown to enhance antitumor immunity, in part by upregulating classical HLA class I molecules [30] and promoting PD-L1/PD-1 checkpoint inhibition [29, 31]. These findings suggest that distinct cannabinoids differentially regulate mechanisms associated with antigen presentation and immune responses, providing a strong rationale for investigating their role as potential immunomodulatory agents in cancer therapy.

Despite growing interest in the immunomodulatory properties of CBD, its potential to regulate classical and non-classical HLA expressions in tumor cells remains poorly understood. To address this gap, we used the JEG-3 human choriocarcinoma cell line, which constitutively expresses HLA-G in addition to the classic HLA class I molecules. This dual expression property provides a robust model to investigate cannabinoid–immune interactions in a single cellular context. In this framework, we examined both the modulation of HLA, classical and non-classical subtypes (focusing particularly on HLA-G) and the involvement of cannabinoid receptors and associated signaling pathways, aiming to elucidate the mechanisms by which CBD may influence tumor–immune dynamics.

## MATERIALS AND METHODS

### Drugs and Chemicals

Cannabidiol (CBD, purity by HPLC: 99.8%), specific inhibitors of Cannabinoids Receptors (CBRs): LY320135 (CB1-selective antagonist), AM630 (CB2-selective antagonist) and CID16020046 (GPR55-specific inhibitor) and BAPTA-AM (cytoplasmic calcium chelator), were used.

The CBD was purchased from Cerilliant^©^ (Texas, USA). It was initially dissolved in dimethyl sulfoxide (DMSO) to a concentration of 250 mM and stored at -20°C. CBD was further diluted with tissue culture medium for *in vitro* studies, keeping the DMSO concentration below 0.2 %. LY320135 (CB1-selective antagonist) and AM630 (CB2-selective antagonist) and CID16020046 (GPR55-selective antagonist) were purchased from Tocris Bioscience (Bristol, UK). BAPTA-AM (cytoplasmic calcium ion chelator) was kindly provided by Dra. Eugenia Alzugaray (Universidad Nacional de La Plata, Argentina)

### Cell culture

Human choriocarcinoma cell line (JEG-3) was used. It was generously provided by Instituto de Fisicoquímica Biológica y Química, Universidad de Bioquímica y Farmacia (UBA-CONICET), Buenos Aires.

This cell line was cultured *in vitro* in Dulbecco’s modified Eagle’s medium (DMEM) supplemented with 10% foetal bovine serum (Gibco^TM^) according to the manufacturer’s protocols and 1% penicillin/streptomycin (GibcoTM) in a 5% CO2, humidified atmosphere at 37 °C until a confluence state of 75%. Cells were regularly dissociated using Trypsine-EDTA 0.25% for further RNA extraction, cDNA synthesis, polymerase chain reaction (PCR) and reverse transcription quantitative polymerase chain reaction (RT-qPCR) analysis.

### Cell viability assay (MTT)

To evaluate the effect of CBD on cell viability, an MTT colorimetric assay was performed using 3-(4,5-dimethyl-2-thiazolyl)-2,5-diphenyl-2H-tetrazolium bromide (Santa Cruz Biotech, USA). JEG-3 cells were seeded at a density of 1 × 10⁴ cells per well in 96-well flat-bottom plates (Corning Inc., USA) in 100 µL of DMEM containing 0.4% DMSO. After a 24-h attachment period, cells were exposed to increasing concentrations of CBD for an additional 24 h. Following treatment, MTT was added at a final concentration of 0.5 mg/mL and incubated for 3 h. The resulting formazan crystals were dissolved by adding 200 µL of 100% DMSO per well, and absorbance was measured at 560 nm with 640 nm as the reference wavelength using an automated microplate reader (Beckman Coulter DTX 880, Fullerton, CA, USA). IC₅₀ values were calculated from concentration–response curves (% cell death) using non-linear regression analysis.

### CBD Treatments with specific receptor inhibitors

Cells were seeded in 6-multi-well plates at a density of 5 × 10⁵ cells/well in complete culture medium and allowed to adhere for 24 h. Following adhesion, five independent assays were performed.

#### Monomeric CBR inhibition assay

Cells were allocated into four experimental groups: (i) negative control (DMEM + 10% FBS + 0.4% DMSO), (ii) positive control (DMEM + 10% FBS + 0.4% DMSO + 1 µM CBD), (iii) inhibitor treatment (DMEM + 10% FBS + 0.4% DMSO + CBR inhibitors), and (iv) inhibitor + CBD treatment (DMEM + 10% FBS + 0.4% DMSO + CBR inhibitors + 1 µM CBD). LY320135, AM630, and CID16020046 were used at a final concentration of 5 µM. Treatments were performed for 24 h, with medium replacement after 12 h.

#### Dimeric CBR inhibition assay

Cells were distributed into the same four groups as described above. Inhibitor treatments consisted of combinations targeting CB1/CB2, CB1/GPR55, or CB2/GPR55, each applied at a final concentration of 5 µM. Treatments were carried out for 24 h, with medium replaced after 12 h of incubation.

After each experiment was completed, samples were collected in TRIzol for RNA extraction.

### Cytoplasmic calcium chelation assay

To assess the potential involvement of intracellular calcium in the signaling pathway underlying CBD-mediated modulation of HLA expression, a cytoplasmic calcium chelation assay was performed. Cells were seeded in 6-multi-well plates at a density of 5 × 10⁵ cells/well in complete culture medium and allowed to adhere for 24 h. Following adhesion, three independent assays were performed. The samples were assigned to four experimental conditions: (i) negative control (DMEM supplemented with 10% FBS and 0.4% DMSO); (ii) positive control (DMEM + 10% FBS + 0.4% DMSO + 1 µM CBD); (iii) BAPTA-only treatments, comprising escalating BAPTA concentrations (0.1, 0.5, and 1 µM); and (iv) BAPTA + CBD treatments, in which cells were pre-incubated for 1 h with BAPTA (0.1, 0.5, or 1 µM) followed by the addition of 1 µM CBD. All experimental conditions were maintained for 24 h, with medium replacement after 12 h.

### RNA Extraction, cDNA Synthesis, and RT-PCR/ RT-qPCR Analysis

For the analysis Cannabinoids Receptors (CBRs), HLA Ia and HLA-G expression in tumor cells cultured under different experimental conditions, reverse transcription-polymerase chain reaction (RT-PCR) was performed using specific primers. RNA extraction from JEG-3 cells, was performed with TRIzol Reagent (Invitrogen). For cDNA synthesis, 500–1000 ng of the total RNA was retro-transcribed with MMLV reverse transcriptase (Promega), according to manufacturer’s instructions.

For the qualitative analysis of gene expression, PCR was performed using the DreamTaq Green PCR Master Mix (2X) (Thermo Scientific, Waltham, MA, USA, cat. no. K1081). Amplicons were subsequently separated by electrophoresis on a 2% agarose gel and visualized by ethidium bromide staining.

For the quantitative analysis of gene expression, RT-qPCR was performed on cDNA samples. The samples were diluted fivefold, and it was performed with StepOne Plus Real Time PCR System (Applied Biosystems). The FastStart Universal SYBR Green Master Mix (Roche) was used for all reactions. Primers efficiency and initial molecule (N_0_) values were determined by LinReg software 3.0, and gene expression was normalized to RPL7 housekeeping gene, for each condition. All the oligonucleotide sequences are listed in Table 1.

**Table 1.**
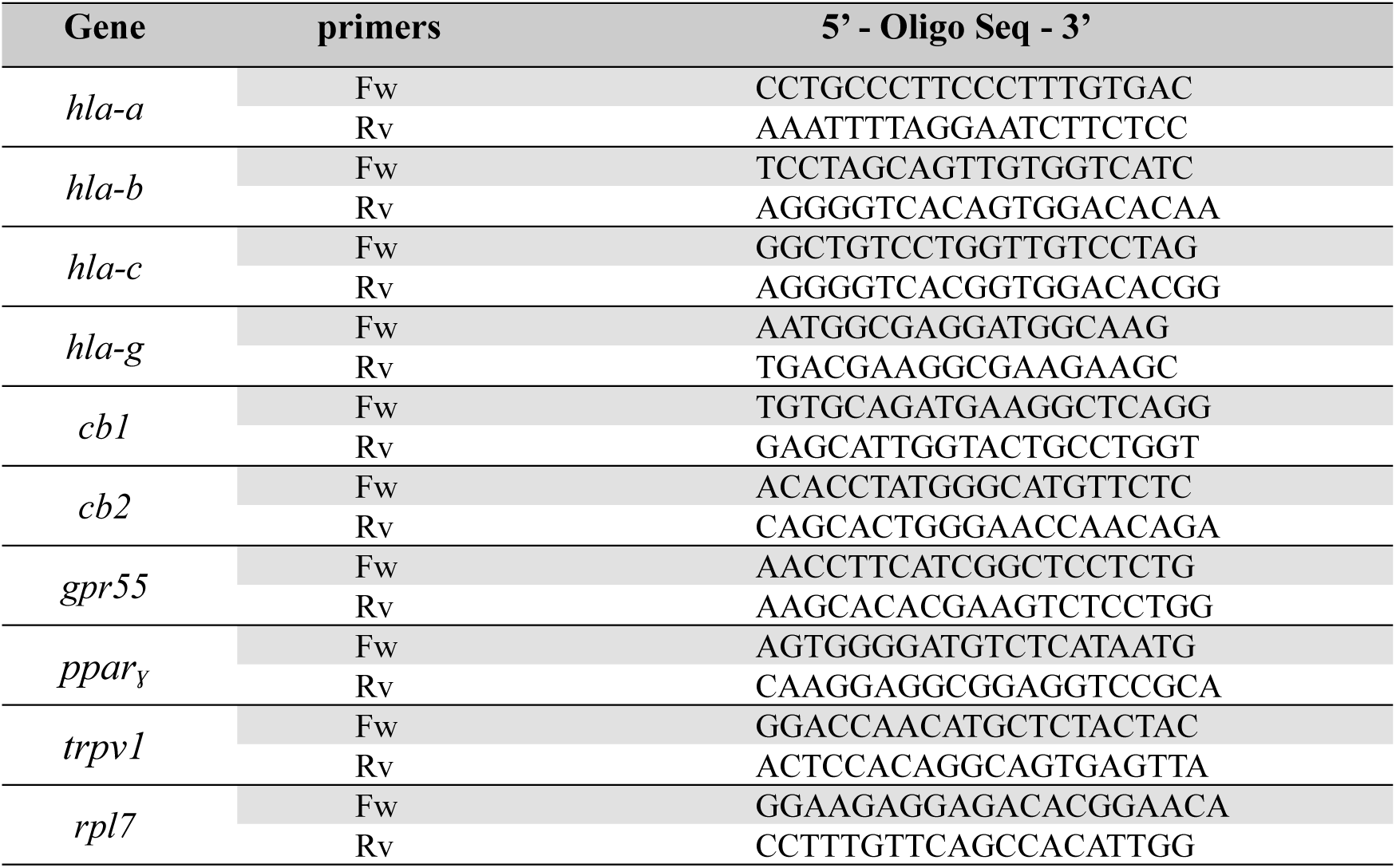
Oligonucleotide sequences of hla classic, *hla-g*, *CBRs*, *pparɣ* and *trpv1* gene and *rpl7* reference gene.

### Statistics

Significant differences were determined using Graph Pad Prism 8 (USA). Statistical significance was calculated using t-tests and ANOVA. Tukey-Kramer post hoc analyses were conducted when appropriate. The significance was set at p < 0.05. Data were expressed as mean ± standard error of mean (SEM).

## RESULTS

### MTT-Based Determination of Non-Cytotoxic CBD Concentrations

To evaluate the potential cytotoxic effect of CBD on JEG-3 choriocarcinoma cells (Figure 1), cells were exposed to increasing concentrations of CBD (0–60 µM). Within the 0–15 µM range, only a slight reduction in viability was detected, with no statistically significant differences among treated groups (p > 0.05). In contrast, concentrations above 15 µM induced an abrupt and significant loss of cell viability (p < 0.001). This assay was essential for establishing sublethal CBD concentrations to ensure that the effects described below reflect physiological cellular responses rather than alternative pathways such as apoptosis. The IC_50_ value obtained was 18.8 µM.

**Figure 1.**
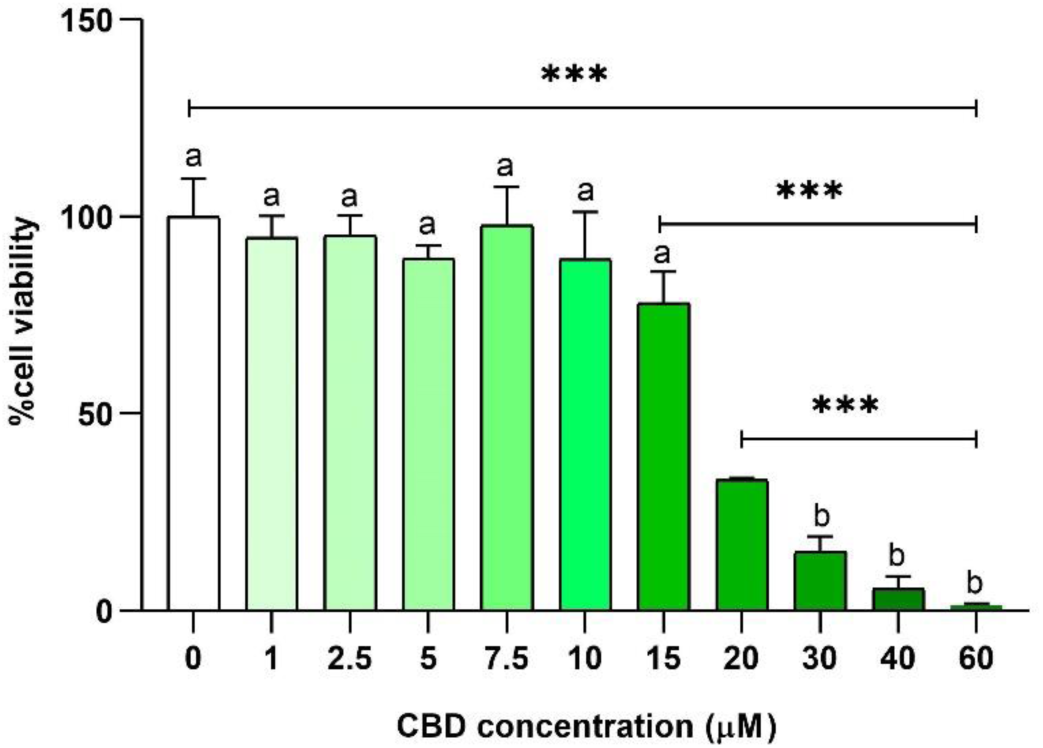
MTT assay. Viability of JEG-3 cells after exposure to increasing CBD concentrations. Bars represent mean ± SEM of eight independent experiments (n = 8). ***p < 0.001. Different letters indicate significant differences.

### Expression profile of cannabinoid-related receptors in JEG-3 cells

To investigate potential signaling pathways regulating HLA-G expression in the JEG-3 choriocarcinoma cell line, we first examined the expression profile of cannabinoid receptors. PCR analysis showed that JEG-3 cells constitutively expressed CB1, CB2 (Figure 2A). In contrast, TRPV1, GPR55 (Figure 2A), and PPARγ (Figure 2B) were not detected under basal conditions. Primer specificity was confirmed using fibroblasts, which showed clear amplification of TRPV1 and GPR55. We next assessed whether exposure to 1 µM CBD modified receptor expression. Under these conditions, the overall expression pattern remained comparable to untreated cells, except for the appearance of a clear GPR55 band (Figure 2A). In addition, PPARγ expression became detectable following CBD treatment (Figure 2B).

**Figure 2.**
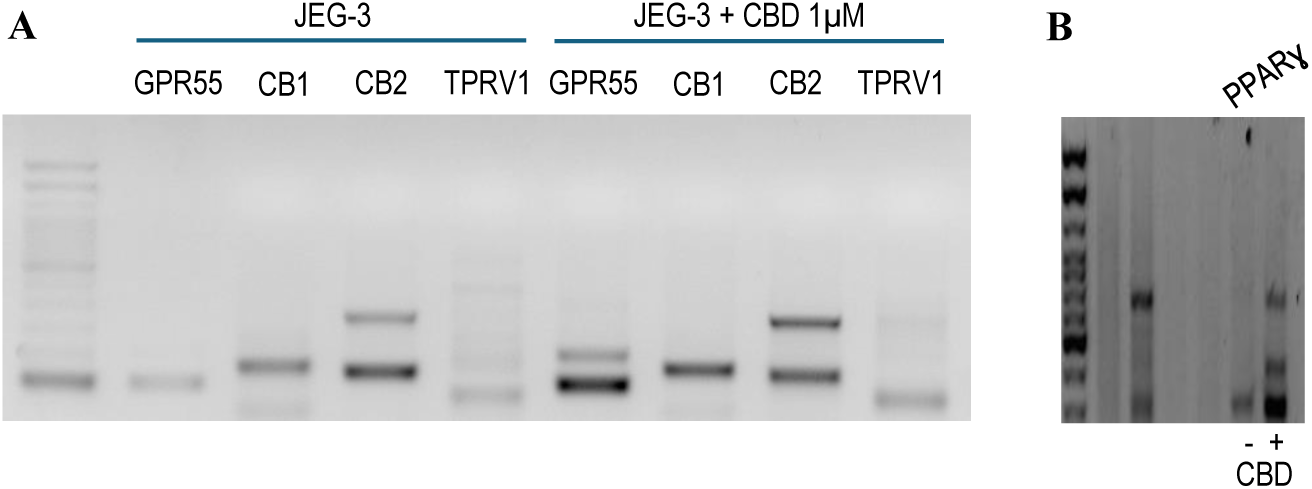
PCR analysis of cannabinoid receptor expression in JEG-3 cells. (A) Agarose gel showing PCR amplification of GPR55, CB1, CB2, and TRPV1. Lane 1: 100 bp ladder; lanes 2–5: control cells; lanes 6–9: cells treated with 1 µM CBD. (B) PCR amplification of PPARγ showing absence of expression in control cells and clear induction following treatment with 1 µM CBD. Five independent biological samples were analyzed for all conditions (n = 5).

These observations were essential for guiding the design of the subsequent cannabinoid receptor inhibition assays.

### CBD upregulates the expression of classical HLA class I molecules in JEG-3 cells

RT-PCR analysis of HLA-A, HLA-B, and HLA-C in JEG-3 cells revealed that treatment with 1 µM CBD induced a marked increase in band intensity for HLA-A and HLA-C, whereas HLA-B showed only a slight elevation compared with controls (Figure 3A). Consistently, RT-qPCR evidenced a significant upregulation (p < 0.001) in classical HLA expression, with locus-specific increases of ∼2.4-fold for HLA-A (Treatment vs. Ct = 2.61 ± 0.08 / 1.08 ± 0.09); 1.5-fold for HLA-B (1.67 ± 0.08 / 0.99 ± 0.09) and 3.4-fold for HLA-C (3.47 ± 0.08 / 1.04 ± 0.07) (Figure 3B).

**Figure 3.**
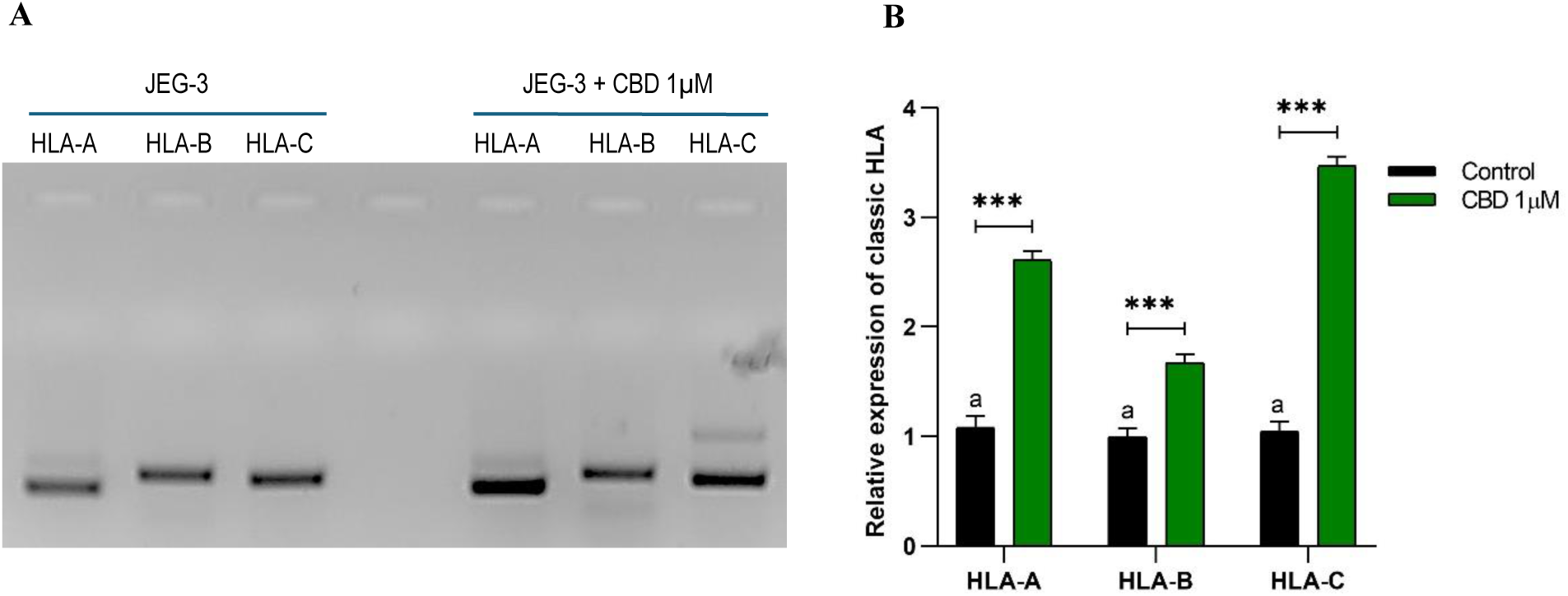
Classical HLA class I expression under CBD treatment in JEG-3 cells. (A) Agarose gel showing PCR products for HLA-A, HLA-B, and HLA-C under control conditions (left) and after treatment with 1 µM CBD (right). Three independent biological samples were analyzed for each condition. (B) Relative expression of classical HLA molecules in control (white bars) and CBD-treated (grey bars) cells. Bars represent the mean ± SEM of three independent experiments (n = 3). ***p < 0.001; the letter “a” indicates no significant differences between groups.

### CBD-induced HLA-C upregulation involves GPR55 and CB2, but not CB1 signaling

To characterize the cannabinoid receptors (CBRs) involved in CBD-mediated regulation of classical HLA molecules, JEG-3 cells were exposed to selective CBR inhibitors in the presence of 1 µM CBD, focusing on HLA-C—the locus exhibiting the highest CBD-induced upregulation. Pharmacological inhibition of CB1 led to a marked increase in band intensity, exceeding that observed with CBD alone. In contrast, CB2 inhibition produced a slight reduction relative to control cells, whereas GPR55 inhibition had no detectable effect (Figure 4A). RT-qPCR analysis corroborated these findings: HLA-C expression significantly increased (p < 0.001) in cells treated with CBD alone (2.9 ± 0.1) or CBD plus CB1 inhibitor (2.7 ± 0.3), whereas CB2 (1.5 ± 0.3) or GPR55 (1.3 ± 0.2) inhibition did not induce significant changes (p > 0.05) relative to baseline (1.1 ± 0.1) (Figure 4C). The potential contribution of CBR dimers to CBD-induced HLA-C upregulation was assessed by RT-qPCR. A significant increase (p < 0.001) of HLA-C was observed in cells treated with CBD (3.0 ± 0.1) or co-treated with CBD plus CB1/CB2 inhibitors (3.0 ± 0.2). In contrast, co-treatment with CBD and CB1/GPR55 (1.2 ± 0.1) or CB2/GPR55 (1.4 ± 0.1) inhibitors did not significantly (p > 0.05) alter expression relative to baseline (1.4 ± 0.1) (Figure 4D).

**Figure 4.**
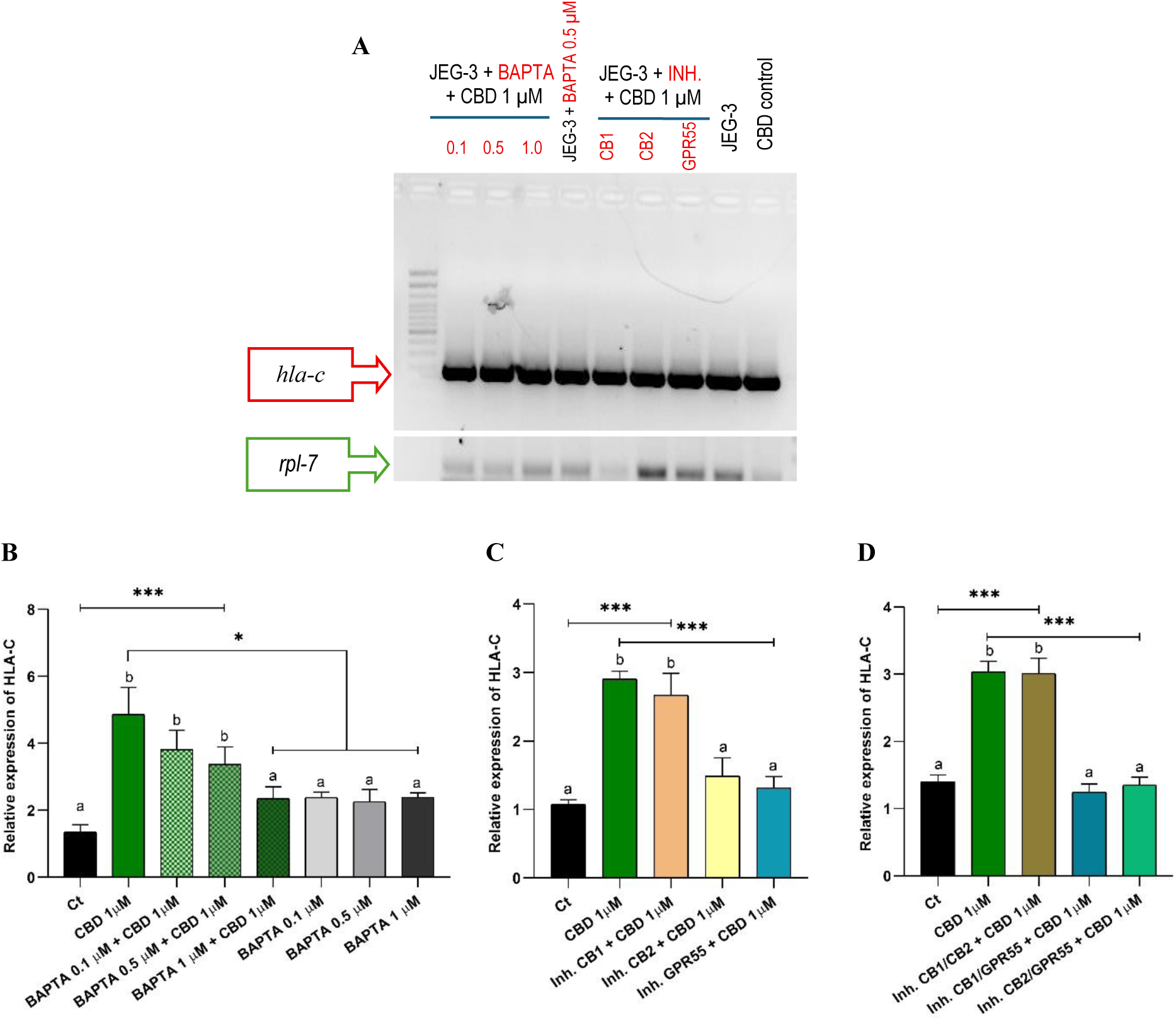
Analysis of calcium chelation and cannabinoid receptor inhibition on HLA-C expression in JEG-3 cells. (A) Agarose gel showing PCR products for HLA-C and RPL7. Lanes 1–4 correspond to calcium ion chelation assays: co-treatments with 1 µM CBD and BAPTA at 0.1, 0.5, and 1 µM, respectively, and cells treated with 0.5 µM BAPTA alone. Lanes 5–7 show cells treated with 1 µM CBD in combination with CB1, CB2, or GPR55 selective inhibitors at 5 µM. Lane 8 corresponds to basal HLA-C expression, and lane 9 to expression after 1 µM CBD. Three independent biological samples were analyzed for each condition. (B) Relative HLA-C expression in cells trated with CBD (1 µM); CBD combined with BAPTA (0.1, 0.5 and 1.0 µM; and BAPTA alone as control (n = 3). (C) Relative HLA-C expression in control cells (without treatment), CBD-treated cells (1 µM), and cells co-treated with CBD (1 µM) combined with CB1, CB2, or GPR55 inhibitors at 5 µM. (D) Relative HLA-C expression in control cells, CBD-treated cells (1 µM), and CBD (1 µM) + receptor inhibitor combinations (CB1/CB2, CB1/GPR55, CB2/GPR55) at 5 µM. In panels C and D, each bar represents mean ± SEM of five independent experiments (n = 5). *p < 0.05 and ***p < 0.001. Different letters indicate significant differences.

### Clasiccal HLA (HLA-C) expression involves calcium signaling

Finally, the role of calcium signaling in HLA-C regulation was examined. RT-PCR band intensities of HLA-C under co-treatment with CBD and increasing BAPTA concentrations suggested that calcium ions are not major contributors, as bands remained above basal levels and comparable to CBD-only treatment (Figure 4A). RT-qPCR analysis showed that co-treatment with CBD and low concentrations of BAPTA (0.1 µM: 3.8 ± 0.6; 0.5 µM: 3.4 ± 0.5) did not differ significantly (p > 0.05) from CBD alone (4.9 ± 0.8). In contrast, increasing BAPTA to 1 µM (2.3 ± 0.3) significantly (p < 0.05) reduced the CBD-induced response, returning expression levels near to baseline. Treatment with BAPTA alone at all tested concentrations (0.1 µM: 2.4 ± 0.2; 0.5 µM: 2.3 ± 0.4; 1.0 µM: 2.4 ± 0.1) produced no changes, yielding values comparable to basal expression. In addition, HLA-C levels in CBD-treated cells, as well as in those exposed to low BAPTA concentrations, remained significantly elevated compared to baseline (1.4 ± 0.2) (p < 0.001) (Figure 4B).

### CB1 receptor inhibition partially restores HLA-G expression

We recently reported that treatment with 1 µM CBD strongly suppresses HLA-G expression in JEG-3 cells [38]. This inhibitory effect clearly contrasts with the CBD-induced upregulation observed for the classical HLA class I molecules described above.

PCR analysis of JEG-3 cells co-treated with 1 µM CBD and 5 µM CBR inhibitors showed that inhibition of CB2 or GPR55 did not modify the suppressive effect of CBD, whereas CB1 inhibition partially restored HLA-G expression (Figure 5A). This effect was validated by RT-qPCR (Figure 5C). No significant differences (p > 0.05) were observed between CBD alone (0.11 ± 0.03) and co-treatments with CBD plus CB2 (0.08 ± 0.04) or GPR55 (0.10 ± 0.02) inhibitors (p > 0.05). In contrast, co-treatment with a CB1 inhibitor (0.42 ± 0.03) significantly increased HLA-G expression compared with CBD alone (p < 0.001), reaching ∼40% of basal levels (1.06 ± 0.14).

**Figure 5.**
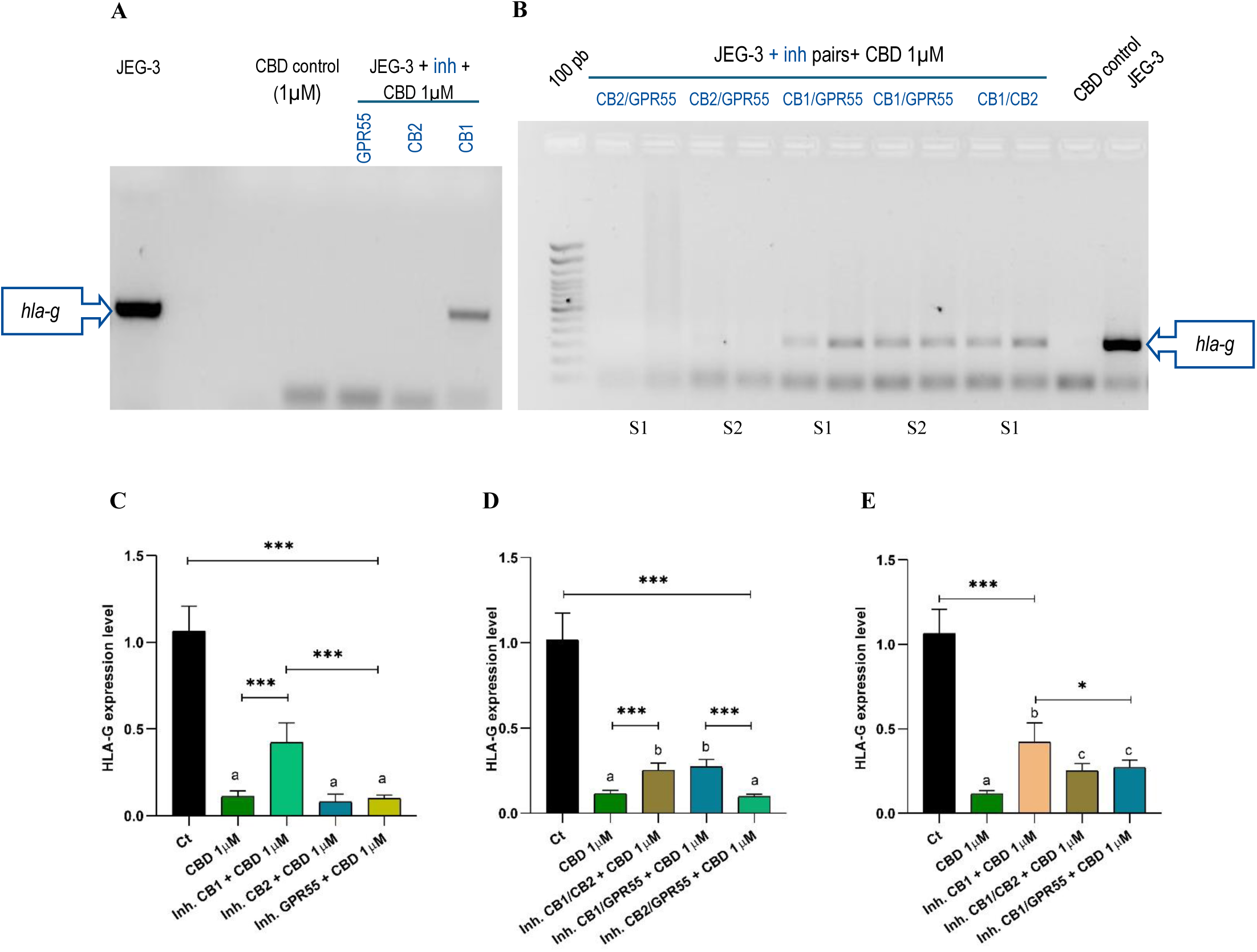
Analysis of HLA-G expression under single or paired CBR inhibition in JEG-3 cells. (A) Agarose gel showing PCR products from the single-receptor inhibition assay. Lane 1: basal HLA-G expression (control); lane 2: HLA-G expression in cells treated with 1 µM CBD; lanes 3–5: HLA-G expression in cells co-treated with 1 µM CBD and the CB1, CB2, or GPR55 inhibitors (5 µM), respectively. (B) Agarose gel showing PCR products corresponding to the paired-receptor inhibition assay. Lane 1: 100 bp ladder; lanes 2–5: HLA-G expression in cells co-treated with 1 µM CBD and CB2/GPR55 inhibitors (samples S1 and S2); lanes 6–9: co-treatment with CBD and CB1/GPR55 inhibitors (S1 and S2); lanes 10–11: co-treatment with CBD and CB1/CB2 inhibitors (S1); lane 12: CBD-treated control (C+); lane 13: untreated control (C–). (C) Relative HLA-G expression for the untreated control, CBD (1µM) treated group, and co-treated groups receiving CBD + single CBR inhibitors (CB1, CB2, or GPR55 at 5 µM). (D) Relative HLA-G expression for the untreated control, CBD (1µM) treated group and co-treated groups receiving CBD + paired CBR inhibitor (CB1/CB2, CB1/GPR55, or CB2/GPR55 at 5 µM). (E) Relative HLA-G expression comparing the untreated control, the CBD group, CBD + CB1 inhibitor and CBD + paired inhibitor involving CB1 (CB1/CB2 and CB1/GPR55). Each bar represents the mean ± SEM of five independent experiments (n = 5). *p < 0.05 and ***p < 0.001. Different letters indicate significant differences.

Because CBRs can form heterodimers upon activation, we next assessed this possibility in CBD-treated JEG-3 cells. Simultaneous inhibition of CB1/CB2 or CB1/GPR55 significantly increased HLA-G expression relative to CBD treatment alone (p < 0.001), whereas inhibition of CB2/GPR55 did not alter expression (Figure 5B). Consistently, RT-qPCR analysis (Figure 5D) showed that co-treatment with CBD and inhibitors of CB1/CB2 (0.21 ± 0.04) or CB1/GPR55 (0.27 ± 0.04) significantly increased HLA-G expression compared with CBD alone (0.11 ± 0.01, p < 0.001), while CBD plus CB2/GPR55 inhibition (0.09 ± 0.01) produced no significant changes (p > 0.05). Figure 5E provides a comparative overview of HLA-G expression after CB1 inhibition alone or combined with CB2 or GPR55 blockade. Although all treatments resulted in significant increases relative to the CBD control, the CB1-independent blockade produced the highest level of HLA-G restoration.

### HLA-G downregulation is dependent on intracellular calcium signaling

In JEG-3 cells co-treated with CBD and the calcium chelator BAPTA, RT-PCR analysis showed a gradual increase in HLA-G amplicon intensity as BAPTA concentrations increased. In contrast, control treatment with BAPTA alone produced no detectable changes (Figure 6A).

**Figure 6.**
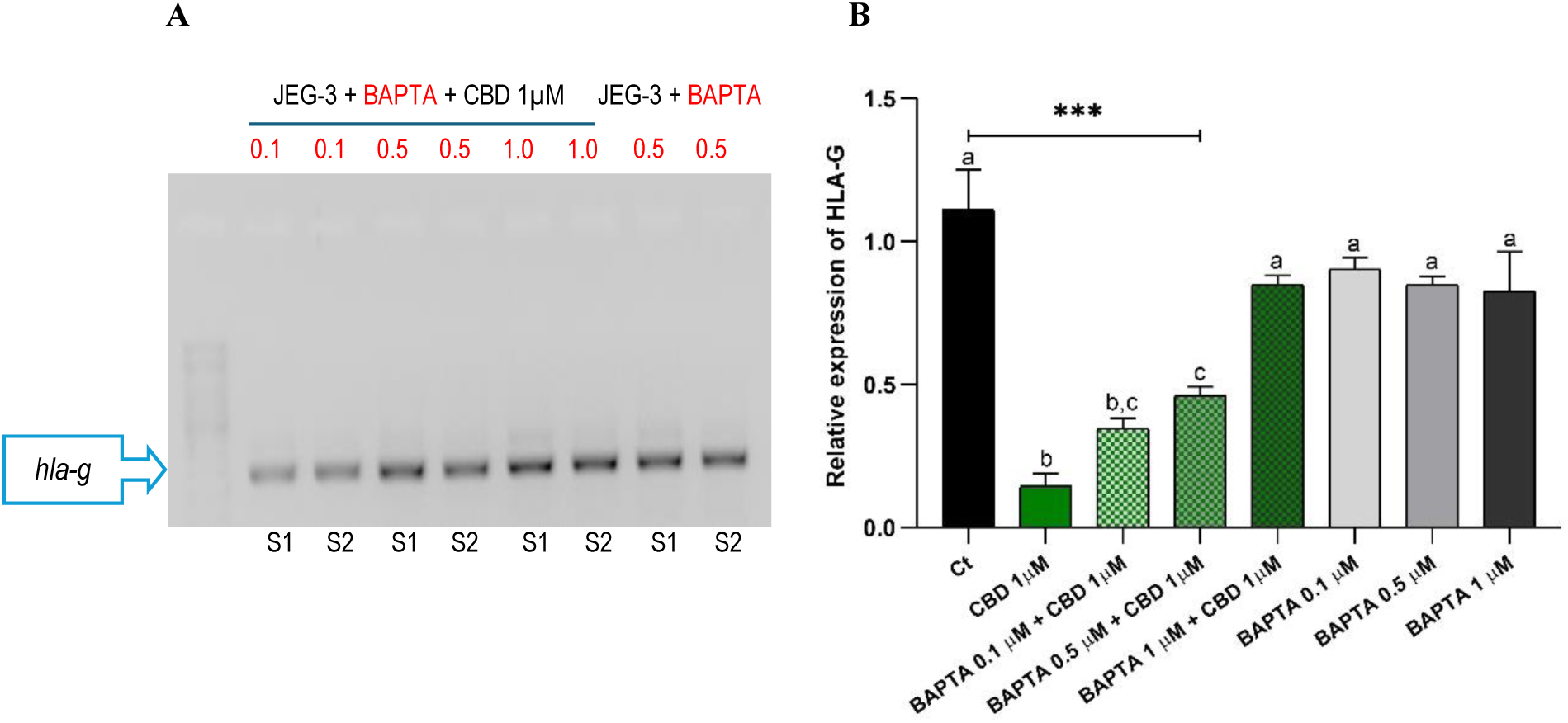
Analysis of calcium-dependent effect in CBD-induced HLA-G expression. (A) Agarose gel showing PCR products from the calcium chelation assay. Lane 1: 100 bp ladder; lanes 2–4: HLA-G expression in JEG-3 cells co-treated with 1 µM CBD and increasing concentrations of BAPTA (0.1, 0.5, and 1 µM); lane 5: HLA-G expression in cells treated with 0.5 µM BAPTA alone. Lanes 6–9: amplification of the reference gene RPL7 corresponding to the conditions described above. (B) Relative HLA-G expression levels for the control group (Ct), cells co-treated with 1 µM CBD and BAPTA at 0.1, 0.5, or 1 µM, and cells treated with BAPTA 0.5 µM alone. Each bar represents the mean ± SEM of three independent experiments (n = 3). ***p < 0.001. Different letters indicate significant differences.

This trend was confirmed by RT-qPCR. Cells treated with 1 µM CBD and increasing concentrations of BAPTA displayed a dose-dependent restoration of HLA-G expression (p<0.001): 0.34 ± 0.04 at 0.1 µM, 0.46 ± 0.03 at 0.5 µM, and 0.85 ± 0.03 at 1 µM, approaching baseline levels. In contrast, treatment with the same concentrations of BAPTA alone (0.1 µM: 0,90 ± 0.04 - 0.5 µM: 0,85 ± 0.03 – 1.0 µM: 0,83 ± 0,10) did not significantly (p>0.05) differ from basal HLA-G expression (1.11 ± 0.20) (Figure 6B).

## DISCUSSION

Tumor immune evasion represents a major challenge for cancer therapy. A comprehensive understanding of the tumor microenvironment and the adaptive mechanisms that enable cancer cells to avoid immune recognition is essential for the design of new and more effective therapeutic strategies. One of the major pathways underlying this process involves dysregulation of HLA expression, which is responsible for antigen presentation in all nucleated cells and represents a key determinant of the immune system’s ability to detect—or overlook—tumor cells. Classical HLA class I molecules (HLA-A, -B, -C) are required for cytotoxic T cell–mediated immune surveillance, whereas non-classical molecules such as HLA-G contribute to immune tolerance and promote tumor escape [32]. Different studies have described that dysregulation of MHC-I expression is associated with tumor progression [33, 34]. Mutations or reduced expression levels of HLA molecules correlate with the development of laryngeal carcinoma and the metastatic spread of melanoma [35, 36] underscoring the central role of HLA profiles in shaping antitumor immunity.

Although the antitumor properties of cannabinoids have been widely explored, their effects on immune regulation remain complex and sometimes contradictory, depending on receptor subtype, tumor context, and ligand specificity [37]. Within this framework, cannabidiol (CBD) has attracted particular interest because of its ability to modulate both tumor cell behavior and immune-regulatory pathways [28–31], with the advantage that—unlike THC—it does not exert psychotropic effects. CBD has been shown to enhance immune-mediated tumor control by upregulating classical HLA molecules in metastatic cancer [30] and reducing PD-1 expression in colon carcinoma [31]. Our previous study provided the first evidence that CBD can downregulate the non-classical HLA-G molecule in JEG-3 choriocarcinoma cells [38], revealing a previously unrecognized connection between cannabinoid signaling and a key immune-evasive pathway. This initial observation raised important questions regarding the specificity of this effect and its potential implications for other HLA class I molecules. In the present work, we expand these findings by incorporating transcriptional analyses of classical HLA class I genes together with receptor-inhibition assays and calcium-modulation experiments, allowing us to dissect the signaling pathways through which CBD regulates both classical and non-classical HLA molecules.

When analyzing the cannabinoid receptor expression profile in JEG-3 cells, several noteworthy observations emerged. Although CB1 and CB2 were constitutively expressed and showed no appreciable differences between control and CBD-treated conditions (1 µM), GPR55 and PPARγ expression became detectable only after CBD exposure. This finding reinforces the relevance of PPARγ as a potential mediator of CBD activity [39, 40]

Our receptor inhibition experiments revealed that CBD-mediated HLA-G reduction is partially reversed by blocking CB1, but not by independently inhibiting CB2 or GPR55 (see figure). Interestingly, we observed that when the selective CB1 inhibitor was combined with either the CB2 or the GPR55 inhibitor, HLA-G expression levels decreased compared to the isolated CB1 inhibitor. However, these values remained significantly higher than the CBD 1 µM control, and higher than those obtained by combining the CB2 and GPR55 inhibitors, which showed expression levels similar to the CBD control. This pattern suggests an absence—or at most a very low contribution—of CB2 and GPR55 in the pathway linking CBD to HLA-G (see Fig. 5).

Taken together, these findings support a selective and predominant involvement of CB1 in CBD-induced HLA-G suppression. While CBD is a weak CB1 agonist, it can also inhibit FAAH and increase anandamide levels, thereby indirectly potentiating CB1 activity [41, 42]. These combined mechanisms may account for the receptor-dependent modulation observed in this study. Consistent with its known biology, the CB2 receptor—predominantly expressed in immune cells—did not contribute to the HLA-G response, aligning with the fact that CB2 is often associated with systemic immunosuppressive pathways of cannabinoids involving classical HLAs [28]. The GPR55 receptor, described mainly in first-trimester fetal endothelial cells [43], also appears to play a minimal role in HLA-G regulation in JEG-3 cells.

In contrast, receptor-inhibition assays performed to dissect the CBD-mediated upregulation of classical HLAs—focusing on HLA-C, given its highest induction—revealed a markedly different pattern. CB1 inhibition did not modify CBD-induced HLA-C expression relative to the CBD control (1 µM). In sharp contrast, independent inhibition of CB2 or GPR55 fully restored HLA-C levels to those of the untreated condition (Fig. 4).

The combination assays further reinforced this interpretation. Co-inhibition of CB1+CB2 produced no reduction relative to CBD alone, whereas inhibition of GPR55—either individually or combined with CB1 or CB2—consistently suppressed the CBD-mediated HLA-C increase back to baseline. CB2 inhibition, both alone and in combination with GPR55, also attenuated the CBD effect, although the contribution of GPR55 appeared more critical within this regulatory pathway. Notably, when CB2 inhibition was combined with CB1 blockade, no reduction was observed relative to CBD treatment, suggesting receptor-specific interactions rather than simple linear additivity.

These findings raise the possibility that receptor dimerization, particularly among CB1, CB2, and GPR55—well-documented in several cellular contexts [37, 44]—contributes to the differential modulation of classical and non-classical HLA molecules. Given that heterodimer function is highly sensitive to cell-type context, receptor stoichiometry, and ligand properties, additional studies using selective agonists and complementary inhibitors will be required to definitively delineate the signaling pathways underlying these responses. Previous studies have proposed a possible functional relationship between PPARγ and GPR55 [45], and both receptors are recognized among the key molecular targets of CBD. However, their interplay in trophoblastic cells remains unclear. Additional experiments will be required to determine whether CBD independently induces the expression of both receptors or whether GPR55 upregulation occurs downstream of PPARγ activation.

Our calcium-modulation experiments indicate that the contribution of intracellular Ca²⁺ to HLA regulation differs markedly between classical and non-classical class I molecules. For HLA-G, chelation of intracellular Ca²⁺ with BAPTA revealed a clear Ca²⁺-dependent mechanism: CBD-induced downregulation was strongly attenuated at 0.1 µM and was almost completely abolished at 1 µM, with expression levels returning close to baseline. In contrast, the response of classical class I HLAs was more attenuated. CBD-induced upregulation of HLA-C showed a modest downward trend at low BAPTA concentrations (0.1–0.5 µM), although these changes did not reach statistical significance. Only at 1 µM BAPTA—in combination with CBD—did HLA-C expression exhibit a significant reduction, returning to levels comparable to the untreated control.

According to our results, the CBD-mediated HLA-G regulatory pathway appears more sensitive to variations in intracellular Ca²⁺, as significant changes were evident even at the lowest BAPTA concentration tested. In contrast, significant modulation of HLA-C required the highest BAPTA dose, suggesting reduced Ca²⁺ dependence.

However, current evidence linking calcium signaling to HLA regulation—particularly HLA-G—remains limited. In the RCC7 HLA-G⁺ clear cell renal carcinoma cell line, the expression of a Ca²⁺ transporter is elevated compared to its HLA-G⁻ counterpart [23]. Yet it remains unclear whether Ca²⁺ influx directly regulates HLA-G expression or whether these changes arise secondarily from HLA-G itself. Within this limited framework, our findings provide new functional support for a Ca²⁺-dependent component in CBD-mediated HLA-G modulation, although further studies will be required to delineate causal relationships and identify the specific signaling pathways involved.

Regarding HLA-C, while an effect is evident, additional experiments will be necessary to clarify its magnitude and mechanism. In particular, determining the source of Ca²⁺ influx—through selective inhibition of plasma membrane and/or endoplasmic reticulum channels—will be essential to fully elucidate the underlying regulatory pathway.

Overall, our findings position CBD as a multifaceted immunomodulatory compound capable of simultaneously reducing tumor immune escape through CB1- and Ca²⁺-dependent suppression of HLA-G, while enhancing immune visibility via GPR55/CB2-mediated upregulation of classical HLA class I molecules. Beyond its previously demonstrated antiproliferative, pro-apoptotic, and anti-migratory effects, this dual immunological action broadens CBD’s relevance in tumor biology. Together, these properties suggest that CBD may not only act as a standalone antitumor agent but could also be compatible with—and potentially supportive of—other immunotherapeutic strategies, raising the prospect that CBD could enhance the effectiveness of existing immune-based treatments.

## ABBREVIATIONS

2-AG: 2 Arachidonoylglycerol
5-HT1A: 5-Hydroxytryptamine
AEA: Anandamide
Ca2+: Calcium ion
CB1: Cannainoid Receptor type 1
CB2: Cannainoid Receptor type 2
CBD: Cannabidiol
CBRs: Cannabinoids Receptors
ECS: Endocannabinoid System
FAAH: Fatty Acid Amide Hydrolase
GABA: Gamma-Aminobutyric Acid,
GPCR: G Protein Coupled Receptors
HLA: Human Leukocyte Antigen
HLA: Ia Classical Human Leukocyte Antigen class I
HLA: Ib Non classical Human Leukocyte Antigen class I
JEG-3: Human choriocarcinoma cell line
MHC: Ia Classical Major Histocompatibility Complex class I
MHC: Ib Non classical Major Histocompatibility Complex class I
MTT: Methyl Thiazolyl Tetrazolium
NK: Natural Killer
PCR: Polymerase Chain Reaction
PD-1: Programmed Death-1
PDAC: Pancreatic Ductal Adenocarcinoma
PD-L1: Programmed Death-Ligand 1
PPAR: Peroxisome Proliferator-Activated Receptor
RCC7: Renal Cell Carcinoma cell line
RT-PCR: Reverse Transcription Polymerase Chain Reaction
THC: Δ9-Tetrahydrocannabinol
TRPV: Transient Receptor Potential Vanilloid

## Acknowledgments

We gratefully acknowledge Dr. M. Eugenia Alzugaray for generously providing the BAPTA reagent for performing the calcium chelation experiments included in this study.

## Author contributions

Project development F.L.R., M.N.G; Experimental designs: F.L.R., K.I.M; Formal analysis: F.L.R., K.I.M., M.B.P., M.N.G., F.J.S.; Methodology: K.I.M, M.B.P.; Resources: F.L.R., M.B.P., M.N.G., F.J.S., E.D.C.; Writing-original draft: F.L.R., K.I.M; Writing-review & editing: F.L.R., K.I.M., M.B.P., M.N.G., F.J.S., E.D.C. All authors contributed to the final draft.

## Corresponding authors

Correspondence to F.L.R. (main corresponding author) or M.N.G.

## Data Availability

The raw data for this study is available from the corresponding author upon reasonable request.

